# A Cre-dependent CRISPR/dCas9 system for gene expression regulation in neurons

**DOI:** 10.1101/2020.11.20.391987

**Authors:** Nancy V. N. Carullo, Jenna E. Hinds, Jasmin S. Revanna, Jennifer J. Tuscher, Allison J. Bauman, Jeremy J. Day

## Abstract

Site-specific genetic and epigenetic targeting of distinct cell populations is a central goal in molecular neuroscience and is crucial to understand the gene regulatory mechanisms that underlie complex phenotypes and behaviors. While recent technological advances have enabled unprecedented control over gene expression, many of these approaches are focused on selected model organisms and/or require labor-intensive customizations for different applications. The simplicity and modularity of CRISPR-based systems have transformed this aspect of genome editing, providing a variety of possible applications and targets. However, there are currently few available tools for cell-selective CRISPR regulation in neurons. Here, we designed, validated, and optimized CRISPR activation (CRISPRa) and CRISPR interference (CRISPRi) systems for Cre recombinase-dependent gene regulation. Unexpectedly, CRISPRa systems based on a traditional double-floxed inverted open reading frame (DIO) strategy exhibited leaky target gene induction in the absence of Cre. Therefore, we developed an intron-containing Cre-dependent CRISPRa system (SVI-DIO-dCas9-VPR) that alleviated leaky gene induction and outperformed the traditional DIO system at endogenous genes in both HEK293T cells and rat primary neuron cultures. Using gene-specific CRISPR sgRNAs, we demonstrate that SVI-DIO-dCas9-VPR can activate highly inducible genes (*GRM2, Tent5b*, and *Fos*) as well as moderately inducible genes (*Sstr2* and *Gadd45b*) in a Cre-specific manner. Furthermore, to illustrate the versatility of this tool, we created a parallel CRISPRi construct that successfully inhibited expression from of a luciferase reporter in HEK293T cells only in the presence of Cre. These results provide a robust framework for Cre-dependent CRISPR-dCas9 approaches across different model systems, and will enable cell-specific targeting when combined with common Cre driver lines or Cre delivery via viral vectors.

## INTRODUCTION

Genetic and epigenetic targeting are fundamental strategies to study gene regulation and function, and also provide novel therapeutic avenues for genetic diseases. In recent years, clustered regularly interspaced short palindromic repeats (CRISPR) systems have revolutionized the field as tools for site-specific DNA and RNA editing (Jinek et al., 2012; Cong et al., 2013; Savell and Day, 2017; Pickar-Oliver and Gersbach, 2019). In these systems, a Cas nuclease is directed to target DNA by an engineered CRISPR single guide RNA (sgRNA), resulting in nuclease-mediated cleavage of the targeted nucleic acid sequence. In addition to classical CRISPR-Cas9 systems, a number of new approaches have been developed based on a mutated, catalytically dead Cas9 (dCas9). dCas9 strategies provide a powerful system to target specific genomic loci without causing double-strand breaks. Instead, dCas9 constructs can be fused to an array of different effector proteins (Savell and Day, 2017). dCas9 systems are commonly used to induce transcriptional activation (CRISPRa) or interference (CRISPRi), epigenetic modifications, chromatin looping, tagging, or can be used as an anchor system to deliver non-coding RNAs (CRISPR-Display) (Hilton et al., 2015; Konermann et al., 2015; Shechner et al., 2015; Yeo et al., 2018; Chen et al., 2019; Savell et al., 2019a; Carullo et al., 2020).

DNA recombinases are commonly used to enable inversion, deletion, or integration of transgenes in a cell-specific manner. ‘The Cre/Lox system is likely the best understood and most commonly used recombination approach in which the Cre recombinase recognizes specific 34bp palindromic Lox sites within a DNA sequence (Anton and Graham, 1995; Gibb et al., 2010). Cre-mediated inversion or deletion occurs when flanking Lox sites are oriented in opposing or parallel orientations, respectively. Importantly, Cre recombinase expression driven under cell type-specific promoters enables targeted gene manipulation of these cell populations (Madisen et al., 2012; Daigle et al., 2018). Due in part to the ease and versatility of this popular system, hundreds of different Cre-driver transgenic lines have been generated for cell-targeted approaches.

Cre-dependent genetic FLEX switches (flip-excision; also known as double-inverted open reading frame, or DIO switches) invert DNA sequences to enable gene activation or silencing (Schnütgen et al., 2003). All FLEX/DIO systems capitalize on Lox sequence variants that provide additional specificity to Cre recognition. Since antiparallel LoxP sites cause continuous inversion, DIO systems use two heterotypic sets of Lox sequences. Recombination at one Lox pair causes transgene inversion, allowing deletion of the genetic material (including a Lox site) between the other Lox pairs. This strategy stops the inversion cycle through the excision of one antiparallel Lox site. A common FLEX/DIO strategy uses LoxP and Lox2272 sequence variants due to their efficient and specific recombination characteristics (Lee and Saito, 1998). This approach has widely been used in neuroscience, enabling cell type-specific optogenetic and chemogenetic activation, overexpression or knockdown of transgenes, and perturbation of endogenous genes (Kühn et al., 1995; Tsien et al., 1996; Gong et al., 2007; Atasoy et al., 2008; Madisen et al., 2012; Erwin et al., 2020).

While CRISPR/dCas9 approaches have enabled targeted induction of transcriptional or epigenetic states at selected genes, inducible cell type-specific CRISPR tools based on these platforms remain limited (Kumar et al., 2018; Bäck et al., 2019). Here, we developed novel Cre-dependent CRISPRa and CRISPRi systems that incorporate a synthetic intron into the dCas9 transgene. Creating an intron-containing dCas9 construct offers two advantages: (1) Introns can increase transgene expression compared to intron lacking counterparts (Shaul, 2017), and (2) Introns provide a non-coding locus for insertion sites of Lox sequences without disturbing the integrity of the coding protein. By separating the dCas9 gene into two segments with a short SV40 intron (SVI), we created CRISPRa and CRISPRi systems in which the first segment of dCas9 is double-floxed and inverted. This approach prevents leaky expression of functional CRISPR constructs by requiring Cre-induced inversion to orient both dCas9 segments into an open reading frame. Furthermore, the construct is designed for easy rearrangement and replacement of promoters and effector fusions and can be applied for various CRISPR/dCas9 systems.

## RESULTS

### Creation and validation of a traditional FLEX-CRISPRa system

AAV-driven FLEX systems are among the most common Cre/Lox approaches used in neurons, enabling inducible expression by combination with Cre-driver animal models. While this strategy has proven to be a versatile approach for cell-specific targeting, few tools have been described for Cre-dependent CRISPR manipulation. Given the flexibility of CRISPRa and CRISPRi systems for transcriptional manipulation at endogenous loci, development of Cre-dependent platforms for these systems are of interest. To create a Cre-inducible system for transcriptional activation, we adapted a CRISPRa tool recently validated in neurons (Savell et al., 2019a) using a common DIO strategy. In this DIO system, an inverted dCas9-VPR cassette was flanked by LoxP and Lox2272 sites (**Figure 1a**), enabling Cre-dependent inversion of dCas9-VPR into the correct orientation followed by the excision of one antiparallel Lox site to prevent further inversion events. To validate this system in dividing cells, HEK293T cells were co-transfected with either a constitutively active dCas9-VPR construct or the DIO-dCas9-VPR construct, in tandem with sgRNA plasmids targeting either the human *GRM2* promoter, or a non-targeting control (sgRNA for the bacterial *lacZ* gene). The DIO-dCas9-VPR construct was tested with or without transfection of a Cre-2A-mCherry plasmid that was driven under the human synapsin (hSYN) promoter (**Figure 1b**). Constitutive activation of the endogenous *GRM2* gene yielded a 75-fold increase of *GRM2* mRNA compared to a non-targeting *lacZ* control sgRNA. While the Cre-dependent DIO-dCas9-VPR system was able to induce *GRM2* mRNA transcription by 112-fold, we also observed a 49-fold increase in the absence of Cre. Next, to examine gene induction in neurons, we transduced rat primary striatal neuron cultures with lentiviruses expressing sgRNAs and CRISPR machinery (**Figure 1c**). CRISPR sgRNAs targeting the *Tent5b* (also known as *Fam46b)* promoter resulted in strong upregulation of *Tent5b* mRNA with both the constitutive and Cre-dependent constructs. However, in neurons the DIO system also exhibited leaky expression, indicated by an 8-fold increase of *Tent5b* mRNA in the absence of Cre. *Tent5b* is a highly inducible gene which could be more susceptible to baseline/leaky induction compared to other genes. Therefore, we tested the DIO-dCas9-VPR system at two other genes including a gene encoding the neuropeptide receptor *Sstr2* and DNA-damage inducible gene *Gadd45b* (**Figure 1d**). Although these genes are less inducible than *Tent5b* (with induction rates of 8-19-fold compared to *lacZ* controls), we still observed non-specific gene induction in the absence of Cre (2-5-fold induction without Cre). This baseline induction of target genes is likely due to leaky expression of the DIO-dCas9-VPR, since no transgene inversion could be detected in PCR verification using genomic DNA isolated from cells lacking Cre (**Figure 1e**). These results are consistent with recent evidence that DIO transgenes can be expressed in the inverted orientation at low levels in the absence of Cre recombinase (Fischer et al., 2019).

**Figure 1.**
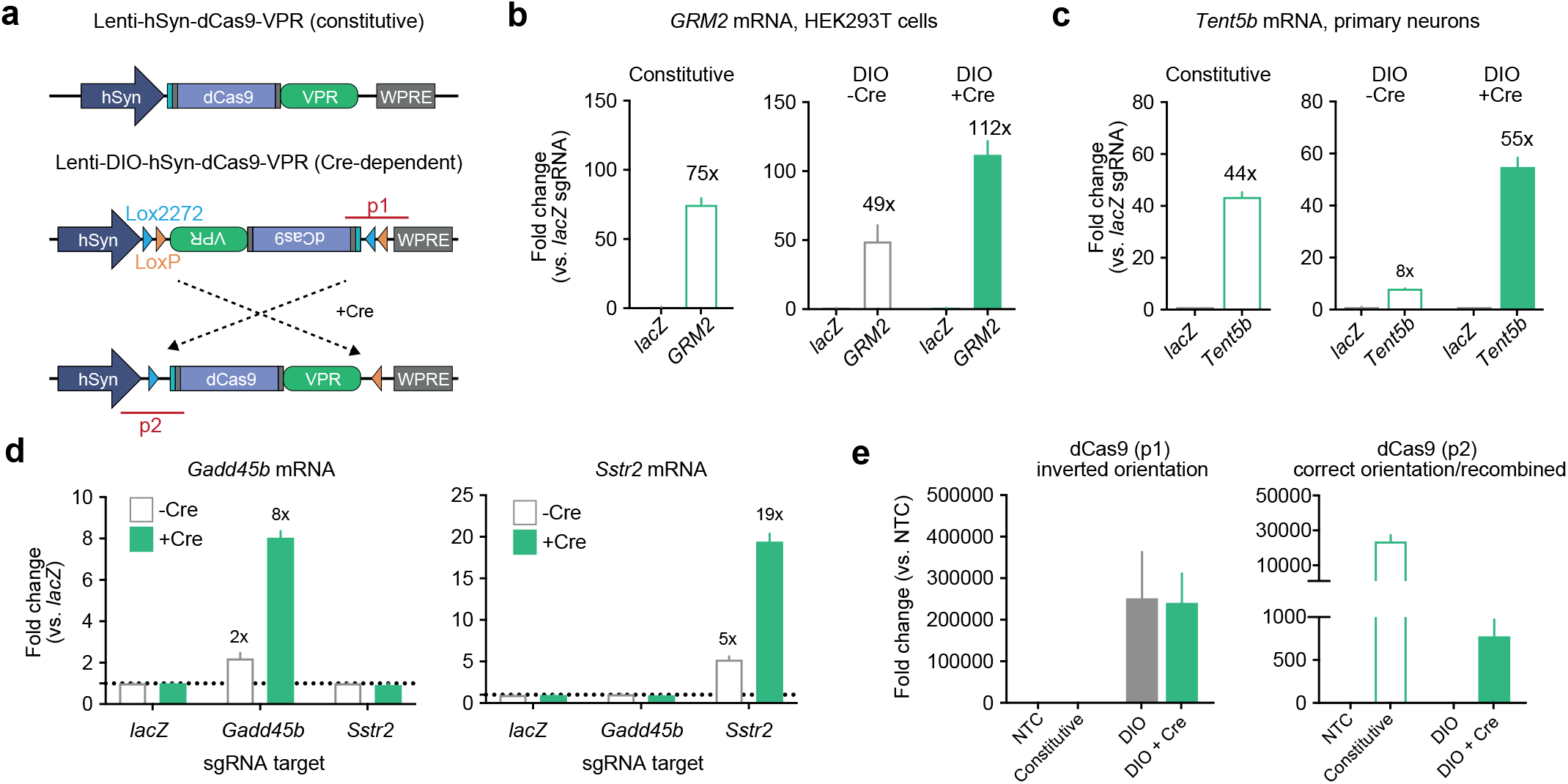
Design and validation of a DIO-dCas9-VPR CRISPRa plasmid. **a**, Illustration of CRISPRa construct designs for a traditional constitutive dCas9-VPR (top) and a Cre-dependent double-floxed inverted open reading frame (DIO) dCas9-VPR (bottom). PCR products for primer sets used in panel e are illustrated as p1 and p2 (red). **b**, In transfected HEK293T cells, the constitutive dCas9-VPR construct induced the endogenous target gene *GRM2* by 75 fold compared to the non-targeting *lacZ* sgRNA control (Welch’s *t*-test *t*(2) = 14.43, *p* = 0.0048). DIO-dCas9-VPR significantly induced transcription of *GRM2* with and without Cre (*n* = 3 per group, two-way ANOVA *F*(1,8) = 32.89, *p* = 0.0004). **c**, Lentiviral expression of constitutive dCas9-VPR construct as well as the DIO-dCas9-VPR construct in rat striatal neurons revealed increased mRNA for the target gene *Tent5b* (Welch’s *t*-test for constitutive *t*(2) = 22.81, *p* = 0.0019, two-way ANOVA for DIO with *n* = 3 per group, *F*(1,8) = 487.9, *p* < 0.0001). **d**, Additional CRISPRa target genes tested in neurons demonstrated leaky induction in the absence of Cre recombinase (two-way ANOVA with *n* = 3 per group for *Gadd45b F*(1,12) = 91.03, *p* < 0.0001, and for *Sstr2 F*(1,12) = 142.1, *p* < 0.0001). **e**, qPCR on transfected HEK293T cell DNA showed high levels of the inverted dCas9-VPR cassette with and without Cre (p1, left, *n* = 4 per group, Kruskal-Wallis test *F*(3,12) = 13.06 *p* < 0.0001). Inversion into the correct orientation only occurred in the presence of Cre (p2, right, n = 4 per group, Kruskal-Wallis test *F*(3,12) = 12.11 *p* = 0.0002). Constitutive VPR and DIO groups (with and without Cre) are compared to a non-transduced control (NTC). Data expressed as mean ± s.e.m.

### Intron insertion into dCas9 increases expression and enables creation of a split-dCas9 FLEX system

Given that an inverted dCas9-VPR cassette still resulted in target gene induction in the absence of Cre, we next aimed to design a system that would prohibit expression of the full length dCas9-VPR fusion protein without Cre-mediated recombination. To achieve this goal, we designed an intron-containing FLEX system. First, we inserted a small SV40 intron (SVI) into the dCas9 sequence to create a constitutively active, intron-containing construct (SVI-dCas9-VPR) in which the dCas9 cassette is divided into two segments by the SV40 intron (**Figure 2a**). We created two versions (1.0 and 2.0) of this SVI-dCas9-VPR construct to determine optimal intron positioning and maximal splicing efficiency. While the intron was only slightly shifted between versions 1.0 and 2.0, SVI-dCas9-VPR 2.0 created a frameshift immediately after the intron, resulting in a premature stop codon within less than 100 bases if the product was not spliced (**Figure 2b**). PCR amplification from cDNA with primers spanning this intron sequence following HEK293T plasmid transfection revealed that both constructs produced detectable dCas9 expression. However, splicing efficiency of the SVI-dCas9-VPR 2.0 version was higher as compared to SVI-dCas9-VPR 1.0 (**Figure 2c, d**).

**Figure 2.**
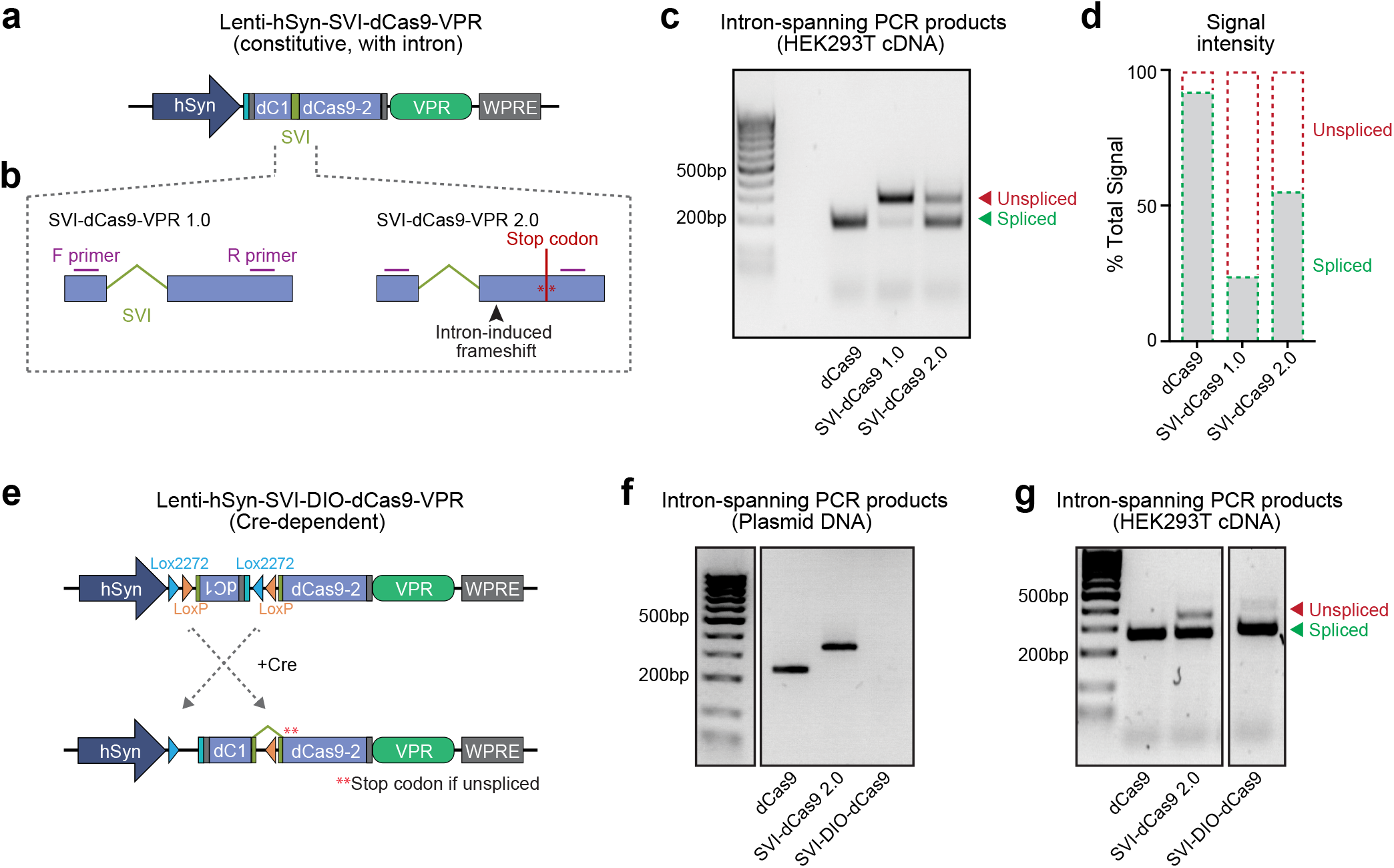
Design of an intron-containing, Cre-dependent CRISPRa system. **a**, Illustration of CRISPR construct design for an intron-containing constitutive dCas9-VPR cassette. **b**, Design of the two different intron positions. **c**, SVI-dCas9-VPR 1.0 and 2.0 intron positions were tested in HEK293T cells. PCR amplification of cDNA with primers spanning the intron within dCas9 showed more efficient splicing of the SVI-dCas9-VPR 2.0 construct. **d**, Quantification of PCR product intensities shown in panel c. **e**, Illustration of a Cre-dependent SVI-DIO-dCas9-VPR construct in which the first dCas9 segment is double-floxed and inverted. **f**, PCR amplification of plasmid DNA with intron-spanning primers verify that no product results from the SVI-DIO-dCas9-VPR plasmid prior to recombination. **g**, PCR amplification of cDNA following HEK293T cell transfection with dCas9-VPR, SVI-dCas9-VPR, and SVI-DIO-dCas9-VPR with Cre showed that SVI-DIO-dCas9-VPR transfected cells express spliced dCas9-VPR transcripts in the presence of Cre recombinase.

Using the more efficient SVI-dCas9-VPR 2.0, we next generated a split-dCas9 DIO cassette by inverting and flanking the first segment of dCas9 with LoxP and Lox2272 sites to prevent leaky transgene expression (**Figure 2e**). With only part of the dCas9 cassette inverted, Cre-dependent inversion and recombination is required to yield a full-length functional dCas9-VPR fusion protein. Using the same intron-spanning primer set as in previous validations, we compared PCR products amplified from the original constitutive dCas9-VPR, the intron-containing constitutive VPR (SVI-dCas9-VPR 2.0), and the new Cre-dependent SVI-dCas9-VPR (SVI-DIO-dCas9-VPR) plasmids. As predicted, the SVI-dCas9-VPR 2.0 plasmid yielded a longer product compared to the original dCas9-VPR with no intron (**Figure 2f**), due to the 97bp intron in the SVI-dCas9-VPR 2.0 plasmid. However, because the SVI-DIO-dCas9-VPR contains an inverted dCas9 segment, the validation primers target the same strand and yield no PCR product. These results further validate the absence of non-specific recombination following plasmid transfection.

We next transfected HEK293T cells with each of these three constructs to compare recombination, expression, and splicing efficiency of the SVI-DIO-dCas9-VPR construct. The SVI-DIO-dCas9-VPR groups also received a Cre-EGFP plasmid that was driven under the hSYN promoter to initiate recombination. PCR amplification of cDNA generated from the transfected HEK293T experiments with intron-spanning primers revealed strong signals for the short, spliced PCR product for all three groups (**Figure 2g**). This indicates that all three constructs were expressed in HEK293T cells and that both SVI-dCas9-VPR 2.0 and SVI-DIO-dCas9-VPR transgenes generated mRNA transcripts with efficient splicing and intron exclusion. Together, these experiments demonstrate that this newly developed intron-containing FLEX system is capable of efficient splicing and transcription and that Cre-dependent recombination is required for inversion and expression of the dCas9-VPR transgene.

### Intron-containing FLEX system drives Cre-dependent CRISPR targeting with no leaky gene induction

Next, we sought to test this newly developed intron-containing FLEX CRISPR system in vitro to validate Cre-specific expression and gene induction (**Figure 3a**). Using a similar experimental design as in the DIO-dCas9-VPR experiments (**Figure 1**), we transfected HEK293T cells with plasmids expressing constitutive dCas9-VPR, SVI-dCas9-VPR, and SVI-DIO-dCas9-VPR constructs, along with sgRNA targeting the *GRM2* gene promoter (or *lacZ* control). Both the intron-containing SVI-dCas9-VPR and constitutive dCas9-VPR caused strong induction of *GRM2* compared to their respective *lacZ* controls (**Figure 3b**). Notably, unlike the classic DIO-dCas9-VPR construct, the SVI-DIO-dCas9-VPR plasmid exhibited very little induction without Cre (2-fold) and strong induction with Cre (66-fold).

**Figure 3.**
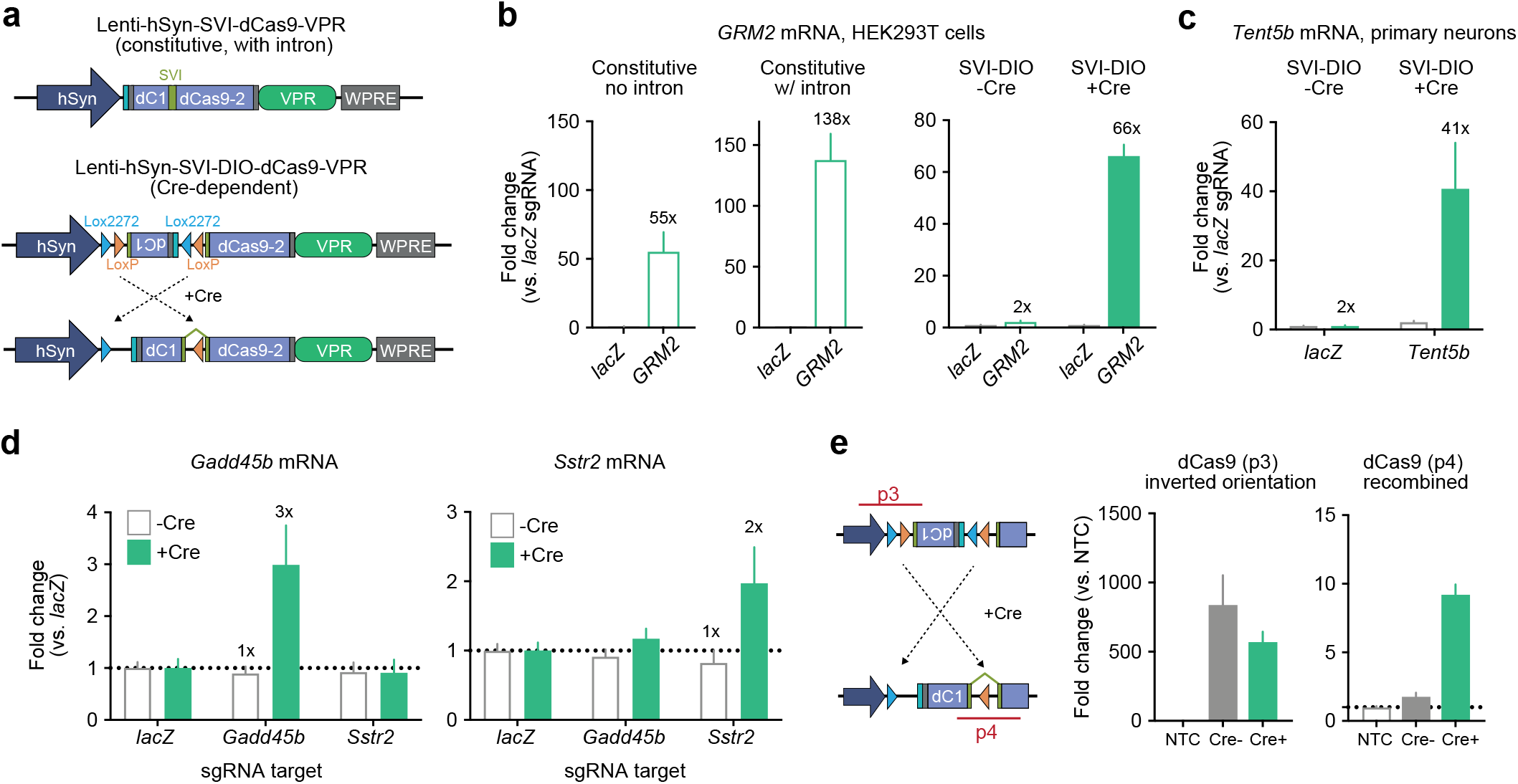
Validation of the SVI-DIO-dCas9-VPR CRISPRa system. **a**, Illustration of CRISPR construct designs for the intron-containing SVI-dCas9-VPR (top) and a Cre-dependent double-floxed inverted open reading frame (DIO) SVI-dCas9-VPR (bottom). **b**, In transfected HEK293T cells the constitutive VPR construct with and without the SV40 intron induced the endogenous target gene *GRM2* by 55 fold and 138 fold compared to the non-targeting *lacZ* control (Welch’s *t*-test for constitutive *t*(2) = 3.846, *p* = 0.0614, and constitutive with intron *t*(2) = 6.322, *p* = 0.0241). SVI-DIO-dCas9-VPR significantly induced transcription of *GRM2* with but not without Cre (*n* = 3 per group, two-way ANOVA *F*(1,8) = 179.5, *p* < 0.0001). **c**, Lentiviral expression of SVI-DIO-dCas9-VPR in striatal neurons increased mRNA for the target gene *Tent5b* (*n* = 10 per group, two-way ANOVA *F*(1,36) = 34.97, *p* < 0.0001). **d**, Additional target genes tested in neurons demonstrated specific induction only in presence of Cre recombinase (two-way ANOVA with *n* = 3 per group for *Gadd45b F*(1, 48) = 4.578, *p* = 0.0375, and for *Sstr2 F*(1,48) = 6.615, *p* = 0.0133). **e**, Illustration of primer and PCR product position for recombination validation experiments (left). qPCR on transduced neuronal DNA revealed high levels of the inverted dCas9-VPR cassette (p3, middle, *n* = 2 per group, Kruskal-Wallis test *F*(2, 4) = 3.714 *p* = 0.2) with and without Cre. Inversion into the correct orientation only occurred in the presence of Cre (p4, right, n = 2 per group, Kruskal-Wallis test *F*(2, 4) = 4.57 *p* = 0.0667). SVI-DIO-dCas9-VPR groups with and without Cre are compared to a non-transduced control (NTC). Data expressed as mean ± s.e.m.

Similarly, targeting the highly inducible *Tent5b* gene in cultured striatal neurons using lentiviral transgene delivery resulted in strong upregulation of *Tent5b* mRNA with the Cre-dependent constructs (41-fold), but not in the absence of Cre (2-fold; **Figure 3c**). These patterns of Cre-dependent induction without leaky effects were also observed for both of the moderately inducible target genes tested in **Figure 1** (*Sstr2* and *Gadd45b*; **Figure 3d**). PCR amplification of genomic DNA that was extracted from transduced primary neurons further validated that recombination required expression of Cre (**Figure 3e**). Together, these data demonstrate that this novel SVI-DIO-dCas9-VPR construct mitigates the leaky gene induction seen in conventional DIO systems for CRISPRa approaches while maintaining the capacity for robust and specific gene induction.

#### Cre-dependent CRISPRa and CRISPRi constructs modulate Fos gene expression at a luciferase reporter

To further validate functionality of our SVI-DIO-dCas9-VPR construct, we tested this system at a luciferase reporter plasmid driven by the rat *Fos* promoter (**Figure 4a, b**) (Duke et al., 2020). Luciferase assays are a common luminescence reporter system used to validate effects of an effector molecules on gene expression. Luciferase, when paired with its substrate D-luciferin and in the presence of ATP, O_2_, and Mg^2+^, produces bioluminescence that can provide insight into the direct transcriptional activity at a regulatory element driving luciferase expression. HEK293T cells were co-transfected with plasmids expressing the *Fos* luciferase plasmid, constitutive or Cre-inducible CRISPRa plasmids, and plasmids expressing sgRNAs targeting either the rat *Fos* promoter or a non-targeting *lacZ* control. Additionally, the SVI-DIO-dCas9-VPR construct was transfected with a Cre-EGFP plasmid. Notably, the constitutive dCas9-VPR constructs with and without the SV40 intron increased luminescence 26- and 25-fold change, respectively, when compared to *lacZ*. Likewise, the SVI-DIO-dCas9-VPR increased luminescence in a similar range (18-fold), but only in the presence of Cre recombinase (**Figure 4c**). Consistent with our previous results in neurons and HEK293T cells (**Figure 3**), this Cre-dependent CRISPRa construct exhibited no leaky induction in the absence of Cre, confirming the utility of this approach for Cre-specific expression and gene induction.

**Figure 4.**
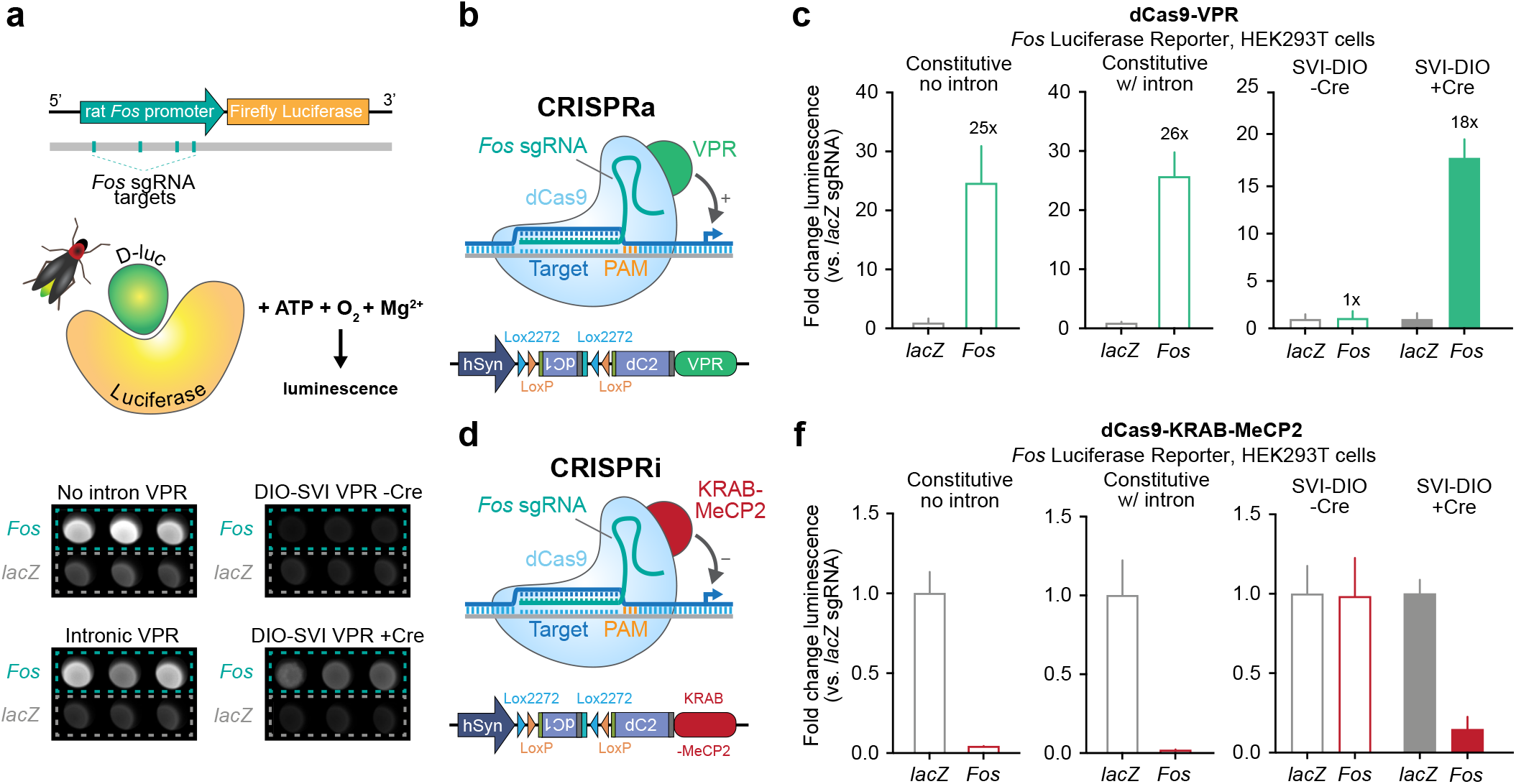
Validation of the SVI-DIO-dCas9-VPR CRISPRa and SVI-DIO-dCas9 CRISPRi systems using a *Fos* luciferase reporter. **a,** Top, illustration of *Fos* Firefly Luciferase plasmid with respective *Fos* sgRNA targets. Bottom, raw luminescence data from VPR luciferase assay illustrates increased luminescence after targeting *Fos* with and without the SV40 intron. SVI-DIO-dCas9-VPR only increases *Fos* luminescence in the presence of Cre. **b,** CRISPRa (VPR) construct coupled with *Fos* Firefly Luciferase plasmid. **c,** In transfected HEK293T cells the constitutive VPR construct with and without the SV40 intron induced a significant increase in luminscence after targeting *Fos* compared to the non-targeting *lacZ* control (Welch’s *t*-test for constitutive VPR no intron *t*(2) = 11.38, *p* < 0.0001, and constitutive with intron *t*(2) = 18.53, *p* < 0.0001). SVI-DIO-dCas9-VPR significantly increased *Fos* luminescence but not without Cre (*n* = 9 per group, two-way ANOVA F(1,32) = 535.4, *p* < 0.0001). **d,** CRISPRi (KRAB-MeCP2) construct coupled with *Fos* Firefly Luciferase plasmid. **f,** After transfection of HEK293T cells, the constitutive KRAB-MeCP2 construct with and without the SV40 intron decreased luminescence when paired with *Fos* sgRNAs (Welch’s *t*-test for constitutive KRAB-MeCP2 no intron *t*(2) = 12.29, *p* = 0.0065, and constitutive with intron *t*(2) = 10.78, p = 0.0001). SVI-DIO-dCas9-KRAB-MeCP2 decreased *Fos* luminescence but not without Cre (*n* = 6 per group, two-way ANOVA F(1,20) = 40.37, *p* < 0.0001). Data expressed as mean ± s.e.m.

While gene overexpression is useful for gain-of-function experiments, CRISPR/dCas9 approaches benefit from the ability to use the same sgRNA for both gene activation and repression, based on the identity of the protein fused to dCas9. In order to generate Cre-dependent system for CRISPRi, we created a parallel Cre-dependent construct in which dCas9 is fused to a KRAB-MeCP2 repressor domain that results in robust gene silencing when targeted to gene promoters (Yeo et al., 2018; Duke et al., 2020) (**Figure 4d**). Following similar transfection and luciferase assay protocols used for our CRISPRi system, we found that constitutive KRAB-MeCP2 both with and without the SV40 intron effectively inhibited luminescence from the *Fos*-luciferase in HEK293T cells. Likewise, the SVI-DIO-dCas9-KRAB-MeCP2 inhibited luciferase activity only in the presence of Cre recombinase. Together, these results demonstrate the capacity of our SVI-DIO CRISPR constructs to target and modulate gene expression at specific transcriptional regulatory regions in a Cre-dependent manner. Likewise, because luciferase reporter assays require successful translation of luciferase protein, these experiments also demonstrate the capability of this system to bidirectionally regulate protein levels in addition to mRNA of targeted genes. Furthermore, adaptation of this SVI-DIO system for multiple CRISPR approaches, such as CRISPRi, highlights the potential and versatility of this tool for cell type- and gene-specific regulation in future experiments.

### DISCUSSION

The mammalian brain consists of heterogeneous cell populations with distinct characteristics and functions. Technologies to study gene function in specific tissues, brain regions, or cell populations are therefore necessary to understand their role in behavior and disease. For example, cell type-selective promoters have been used for specific targeting of excitatory neurons (*Camk2a*) (Mima et al., 2001), astrocytes (*Gfap*) (Morelli et al., 1999), and cell populations that express specific enzymes or receptors such as tyrosine hydroxylase (*Th*) (Savitt et al., 2005), dopamine or serotonin transporters (*Slc6a3, Slc6a4*) (Zhuang et al., 2005; Bäck et al., 2019), or dopamine receptors (*Drd1, Drd2*) (Lobo et al., 2006; Zhang et al., 2006). In this study we developed and optimized Cre-dependent CRISPRa and CRISPRi systems that enable gene and cell type-specific transcriptional activation and inactivation, respectively. A traditional Cre/Lox approach in which the entire dCas9-VPR cassette was double-floxed and inverted was prone to leaky target gene induction in the absence of Cre. While we don’t know with certainty what drives this leaky expression, it is conceivable and in line with recent findings that DIO transgenes can be expressed in reversed orientation in the absence of Cre (Fischer et al., 2019). To avoid this leaky expression, we generated a novel SVI-DIO-dCas9 system in which dCas9 was broken up into two segments by an SV40 intron. This intron provided a location for insertion of Lox sites to invert and double-flox the first dCas9 fragment while leaving the second fragment undisturbed. Without Cre-mediated recombination, the dCas9 segments remain in opposing orientations and are thus unable to yield functional dCas9 protein. In cultured dividing and non-diving cell lines, we demonstrate that the SVI-DIO-dCas9 based CRISPRa system alleviates the leaky transcriptional induction detected in DIO CRISPRa systems (**Figure 1, Figure 3**). This work provides evidence that the SVI-DIO-dCas9-VPR construct can activate highly inducible endogenous genes in HEK293T cells (*GRM2* gene) and striatal neurons (*Tent5b*), without undesired gene induction in the absence of Cre recombinase (**Figure 3, Figure 4**). This system also performed well when targeted to moderately inducible genes such as *Sstr2* and *Gadd45b* in neurons (**Figure 3**), demonstrating a wide range of effect size and application possibilities. Furthermore, successful activation and repression of a *Fos* luciferase reporter in HEK293T cells using SVI-DIO CRISPR constructs demonstrates the versatility of this machinery to provide robust bidirectional control over gene expression and protein levels.

One of the first cell type-specific CRISPR systems used in neurons was based on Cre-dependent expression of CRISPR sgRNAs. Bäck et al. developed a Lox-stop-Lox based sgRNA construct and used it for CRISPR-Cas9 mediated gene knock out (Bäck et al., 2019). This study tested this approach in transgenic rats that expressed Cre recombinase under the rat dopamine transporter (*Slc6a3*) promoter to specifically target and knock out the tyrosine hydroxylase (*Th*) gene in dopaminergic neurons. While this approach allowed for cell type-specific expression of sgRNA constructs, a concern with similar systems is the untargeted overexpression of Cas9 or dCas9 fusion proteins, which could potentially cause unintended and non-specific effects on gene expression.

Our system extends this previous work in two ways. First, the SVI-DIO-dCas9 approach alleviates some of these concerns, as the dCas9 construct itself is Cre-dependent, and therefore, functional fusion proteins are not expressed without Cre recombinase. Secondly, our SVI-DIO-dCas9 system is compatible with traditional validated sgRNA constructs, which can be multiplexed for effect size titration at a single gene (Savell et al., 2019a). Additionally, this approach is also compatible with more complex sgRNA arrays that target entire gene programs (Savell et al., 2020). Thus, this study adds another approach for gene regulation to the CRISPR toolbox, while also outlining a novel intron/DIO strategy to avoid leaky Cre-independent transgene expression. Compared to traditional Cre-dependent overexpression vectors, Cre-dependent CRISPR strategies provides several advantages as they enable targeting of one or multiple endogenous genomic loci and titration of effect size for more physiological expression levels.

Prior work has incorporated temporal specificity into Cre/Lox systems, either via use of optically or chemically inducible proteins. For example, selective estrogen receptor modulator (SERM) inducible systems have been generated by fusion of the ligand binding domain of the estrogen receptor to Cre (Cre-ER). In this approach, a mutated version of the mouse ER that binds tamoxifen but not estrogen was used to create a Tamoxifen-inducible Cre system (Danielian et al., 1993; Metzger and Chambon, 2001). Activity of Cre recombinase can therefore be activated or inactivated upon Tamoxifen injections, increasing temporal control of gene editing events.

In addition to cell type-specificity, selective promoters can be used to drive expression in response to stimulation or experience. Bacterial artificial chromosome (BAC) transgenic animals were generated to express GFP or channelrhodopsin (ChR2) under an immediate early gene promoter such as *Fos* or *Arc* (Guenthner et al., 2013). Using activity-induced approaches in combination with our SVI-DIO-dCas9 system would allow for experience-dependent genetic and epigenetic manipulations.

Cre/Lox systems are also among the most common approaches for intersectional circuit-specific manipulations in neuroscience. For example, in recent work, circuit-specific CRISPR genome editing was achieved by injection of a Cre-dependent Cas9 viral vector in the nucleus accumbens (NAc) and injection of a monosynaptic rabies virus expressing a sgRNA targeting the *Fosb* gene in the ventral hippocampus (vHPC) (Eagle et al., 2020). While the sgRNA virus successfully spread through all hippocampal projections, significant knockout of *Fosb* occurred only in vHPC neurons that projected to the NAc. In future studies, our SVI-DIO-dCas9 system could similarly be used to gain a better understanding of projections between specific cell types in the brain. The intron-containing effector construct could be delivered to one brain region and a Cre-expressing construct to the second brain region through a monosynaptic rabies virus or retrograde AAV serotype. Genetic or epigenetic editing would then only occur in cells with projections between the two targeted brain regions without leaky expression or side effects related to effector protein overexpression in non-targeted cells.

While the use of our SVI-DIO-dCas9 system in Cre driver animal models is an expected application, this approach does not require transgenic organisms. In addition to sgRNA and SVI-DIO-dCas9-VPR or SVI-dCas9-KRAB-MeCP2 delivery, a separate construct expressing Cre can be delivered as necessary to target brain tissues. Additionally, this intron-containing dCas9 provides a basic framework that can be customized and combined with a number of effector proteins to introduce a variety of genetic or epigenetic modifications. The system can easily be customized for expression in various cell types and brain regions and adjusted for inducible efforts and enhanced temporal specificity.

### MATERIALS AND METHODS

#### Cultured neuron experiments

Primary rat neuronal cultures were generated from embryonic day 18 rat striatal tissue as described previously (Savell et al., 2019a). Briefly, cell culture wells were coated overnight at 37° C with poly-L-lysine (0.05 mg/ml for culture wells supplemented with up to 0.05 mg/ml Laminin) and rinsed with diH2O. Dissected tissues were incubated with papain for 25 min at 37°C. After rinsing in Hank’s Balanced Salt Solution (HBSS), a single cell suspension of the tissue was re-suspended in Neurobasal media (Invitrogen) by trituration through a series of large to small fire-polished Pasteur pipets. Primary neuronal cells were passed through a 100 μM cell strainer, spun and re-suspended in fresh media. Cells were then counted and plated to a density of 125,000 cells per well on 24-well culture plate with or without glass coverslips (60,000 cells/cm). Cells were grown in Neurobasal media plus B-27 and L-glutamine supplement (complete Neurobasal media) for 11 DIV in a humidified CO2 (5%) incubator at 37° C.

For viral transduction, cells were transduced with lentiviruses on DIV 4 or 5. All viruses had a minimum titer of 1×10^9^ GC/ml, with a target multiplicity of infection (MOIs) of at least 1000. After an 8-16 hr incubation period, virus-containing media was replaced with conditioned media to minimize toxicity. A regular half-media change followed on DIV 8. On DIV 11, transduced cells were imaged and virus expression was verified prior to RNA extraction. EGFP and mCherry expression was also used to visualize successful transduction using a Nikon TiS inverted epifluorescence microscope.

#### RNA extraction and RT-qPCR

Total RNA was extracted (RNAeasy kit, Qiagen) with DNase treatment (RNase free DNAse, Qiagen), and reverse-transcribed (iScript cDNA Synthesis Kit, Bio-Rad). cDNA was subject to qPCR for genes of interest, as described previously (Savell et al., 2016). A list of PCR primer sequences is provided in **Supplementary Data Table 1**.

#### CRISPR-dCas9 construct design

To achieve transcriptional activation or inactivation, lentivirus-compatible plasmids were engineered to express dCas9 fused to either VPR or KRAB-MeCP2, based on existing published plasmids (Addgene plasmid # 114196 (Savell et al., 2019a); Addgene plasmid # 155365 (Duke et al., 2020)). A Cre-dependent DIO version of the dCas9-VPR construct was generated by insertion of LoxP and Lox2272 sequences flanking the dCas9-VPR cassette. dCas9-VPR was PCR-amplified to insert additional restriction sites (KpnI, BmtI, and BspDI) to allow for LoxP and Lox2272 insertion and was subsequently inserted in reverse orientation to create an intermediate construct via sequential digest and ligation (AgeI, EcoRI). LoxP and Lox2272 sites were amplified from a DIO construct (Addgene plasmid # 113685 (Don et al., 2017)) to create restriction sites (KpnI and BmtI around one set; BspDI and KpnI around another set) and were inserted into the intermediate construct via sequential restriction digest and ligation (KpnI, BmtI, BspDI and EcoRI). Intron-containing dCas9-VPR was constructed by the insertion of a gBlock containing the SV40 intron into the original dCas9-VPR plasmid via Gibson assembly (Gibson Assembly kit, New England BioLabs). The SVI-FLEX construct was built via Gibson assembly of the original dCas9-VPR backbone and two gBlocks encoding the SV40 intron sequence, LoxP and Lox2272 sites (sequences based on DIO construct described above) and dCas9-part1 in inverted orientation. The intron-containing CRISPRi construct (SVI-dCas9-KRAB-MeCP2) was built via restriction digest and Gibson assembly (SfiI, EcoRI, and Xhol, Gibson Assembly kit, New England BioLabs) of the KRAB-MeCP2 and SVI-dCas9-VPR constructs. A Cre-encoding construct (Addgene plasmid # 49056 ((Kaspar et al., 2002)) was used to amplify and insert a Cre transgene into a lentivirus compatible backbone that contained the hSYN promoter and expressed mCherry for visualization via Gibson assembly. To create an additional Cre construct, mCherry was replaced with GFP via sequential digest and ligation (EcoRI and XhoI). dCas9-VPR-expressing constructs were co-transduced with sgRNA-containing constructs. Gene-specific sgRNAs were designed using an online sgRNA tool, provided by the Zhang Lab at MIT (crispr.mit. edu) and inserted in a previously described lentivirus compatible sgRNA scaffold construct (Addgene plasmid # 114199 (Savell et al., 2019b)). To ensure specificity, all CRISPR crRNA sequences were analyzed with the National Center for Biotechnology Information’s (NCBI) Basic Local Alignment Search Tool (BLAST) and Cas-OFFinder (http://www.rgenome.net/cas-offinder/). sgRNAs were designed to target *GRM2*, *Tent5b, Sstr2*, *Gadd45b*, and *Fos* respectively. A list of the target sequences is provided in **Supplementary Data Table 1**. crRNA sequences were annealed and ligated into the sgRNA scaffold using the BbsI or BsmBI cut site. Plasmids were sequence-verified with Sanger sequencing; final crRNA insertion was verified using PCR. Lentivirus-compatible SYN-SVI-DIO-dCas9-VPR and SYN-SVI-DIO-dCas9-KRAB-MeCP2 plasmids will be made available on Addgene (plasmid #164576 and # 170378).

#### Lentivirus production

Viruses were produced in a sterile environment subject to BSL-2 safety by transfecting HEK293T cells with specified CRISPR-dCas9 plasmids, the psPAX2 packaging plasmid, and the pCMV-VSV-G envelope plasmid (Addgene plasmids #12260 & #8454) with FuGene HD (Promega) for 40-48 hrs as previously described (Savell et al., 2019b). Viruses were purified using filter (0.45 μm) and ultracentrifugation (25,000 rpm, 1 hr 45 min) steps. Viral titer was determined using a qPCR Lentivirus Titration Kit (Lenti-X, qRT-PCR Titration Kit, Takara). For smaller scale virus preparation, each sgRNA plasmid was transfected in a 12-well culture plate as described above. After 40-48 hr, lentiviruses were concentrated with Lenti-X concentrator (Takara), resuspended in sterile PBS, and used immediately. Viruses were stored in sterile PBS at −80°C in single-use aliquots.

#### HEK293T cell culturing and transfection

HEK293T cells were obtained from American Type Culture Collection (ATCC catalog# CRL-3216, RRID:CVCL_0063) and cultured in standard HEK Media: DMEM (DMEM High glucose, pyruvate; Gibco 11995081) supplemented with 10% bovine serum (Qualified US Origin; BioFluid 200-500-Q) and 1U Penicillin-Streptomycin (Gibco 15140122). Cells were maintained in T75 or T225 tissue flasks. At each passage, cells were trypsinized for 1-3 min (0.25% trypsin and 1 mM EDTA in PBS pH 7.4) at room temperature. For transfection experiment cells were plated in 24 well plates and transfected with FuGene HD (Promega).

#### Luciferase assay

Bidirectional regulation by SVI-DIO CRISPRa and CRISPRi machinery was examined using a previously described *Fos* luciferase reporter plasmid (Duke et al., 2020). 80,000 HEK293T cells were plated in 500 μl HEK Media. After cells reached 40-50% confluence, 500 ng total plasmid DNA was transfected with 1.5 μl FuGene HD (Promega) as follows: 50 ng of luciferase plasmid, 450 ng in 1:2 molar ratio of total sgRNA:CRISPRa or CRISPRi plasmid. A luciferase glow assay was performed according to manufacturer’s instructions 40 hrs following transfection (Thermo Scientific Pierce Firefly Glow Assay; Thermo Scientific 16177). Cells were lysed in 100 μl 1x Luciferase Cell Lysis Buffer while shaking at low speed and protected from light for 30 min. 20 μl of cell lysate was then added to an opaque 96-well microplate (Corning 353296) and combined with 50 μl 1x D-Luciferin Working Solution supplemented with 1x Firefly Signal Enhancer (Thermo Scientific Pierce Firefly Signal Enhancer; Thermo Scientific 16180). Following a 10 min dark incubation period to allow for signal stabilization, luminescence was recorded using a Synergy 2 Multi-Detection Microplate Reader (BioTek). Luminescence in dCas9-KRAB-MeCP2 and dCas9-VPR experiments was recorded with a read height of 1 mm, 1 s integration time, and 135 or 100 ms delay, respectively. Representative images of luciferase reporter activity presented in **Figure 4** were captured using an Azure c600 imager (Azure Biosystems).

## ACKNOWLEDGEMENTS

We thank all current and former Day Lab members for assistance and support. This work was supported by NIH grants MH114990, DA039650, and DA034681 and the UAB Pittman Scholar Program (JJD), and the CIRC Emerging Scholar Award (NVNC).

## AUTHOR CONTRIBUTIONS

N.V.N.C., J.E.H., J.S.R. and J.J.D. conceived of and performed experiments. N.V.N.C., J.E.H., and J.J.D. wrote the manuscript. J.S.R. designed and built the traditional DIO-VPR construct. N.V.N.C. designed and built SVI-VPR, SVI-DIO-VPR, and SVI-DIO-KRAB-MeCP2 constructs with assistance from J.J.T. and A.J.B.. HEK cell transfections, RNA processing were conducted by J.S.R. and N.V.N.C.. Luciferase assays and respective HEK293T cell transfections were performed by J.E.H.. DNA and cDNA PCR-verifications were conducted by N.V.N.C. Lentivirus production, and transductions were performed by J.S.R. and N.V.N.C. with help and assistance from J.J.T.. Primary neuronal cell cultures were generated by A.J.B., J.J.T., and N.V.N.C.. Data was analyzed by N.V.N.C. and J.E.H.. J.J.D. supervised all work.

---

**Supplemental Data Table 1.**
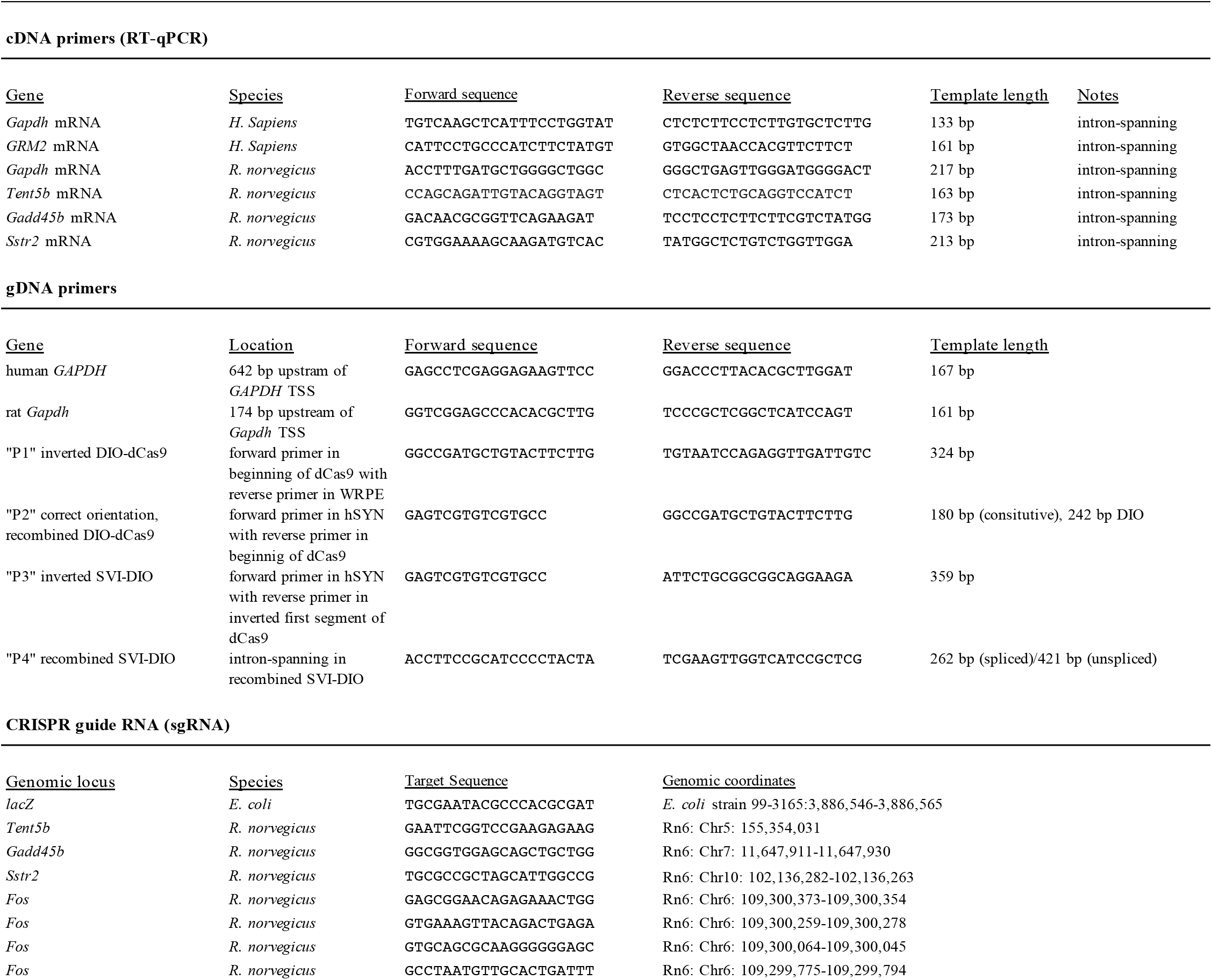
Sequences of primers and sgRNAs

## REFERENCES

Anton M, Graham FL (1995) Site-specific recombination mediated by an adenovirus vector expressing the Cre recombinase protein: a molecular switch for control of gene expression. J Virol 69:4600–4606.

Atasoy D, Aponte Y, Su HH, Sternson SM (2008) A FLEX switch targets Channelrhodopsin-2 to multiple cell types for imaging and long-range circuit mapping. J Neurosci 28:7025–7030.

Bäck S et al. (2019) Neuron-Specific Genome Modification in the Adult Rat Brain Using CRISPR-Cas9 Transgenic Rats. Neuron 102:105–119.e8.

Carullo NVN, Phillips Iii RA, Simon RC, Soto SAR, Hinds JE, Salisbury AJ, Revanna JS, Bunner KD, Ianov L, Sultan FA, Savell KE, Gersbach CA, Day JJ (2020) Enhancer RNAs predict enhancer-gene regulatory links and are critical for enhancer function in neuronal systems. Nucleic Acids Res 48:9550–9570.

Chen L-F, Lin YT, Gallegos DA, Hazlett MF, Gómez-Schiavon M, Yang MG, Kalmeta B, Zhou AS, Holtzman L, Gersbach CA, Grandl J, Buchler NE, West AE (2019) Enhancer Histone Acetylation Modulates Transcriptional Bursting Dynamics of Neuronal Activity-Inducible Genes. Cell Rep 26:1174–1188.e5.

Cong L, Ran FA, Cox D, Lin S, Barretto R, Habib N, Hsu PD, Wu X, Jiang W, Marraffini LA, Zhang F (2013) Multiplex genome engineering using CRISPR/Cas systems. Science 339:819–823.

Daigle TL et al. (2018) A Suite of Transgenic Driver and Reporter Mouse Lines with Enhanced Brain-Cell-Type Targeting and Functionality. Cell 174:465–480.e22.

Danielian PS, White R, Hoare SA, Fawell SE, Parker MG (1993) Identification of residues in the estrogen receptor that confer differential sensitivity to estrogen and hydroxytamoxifen. Mol Endocrinol 7:232–240.

Don EK, Formella I, Badrock AP, Hall TE, Morsch M, Hortle E, Hogan A, Chow S, Gwee SSL, Stoddart JJ, Nicholson G, Chung R, Cole NJ (2017) A Tol2 Gateway-Compatible Toolbox for the Study of the Nervous System and Neurodegenerative Disease. Zebrafish 14:69–72.

Duke CG, Bach SV, Revanna JS, Sultan FA, Southern NT, Davis MN, Carullo NVN, Bauman AJ, Phillips RA, Day JJ (2020) An improved crispr/dcas9 interference tool for neuronal gene suppression. Front Genome Ed 2.

Eagle AL, Manning CE, Williams ES, Bastle RM, Gajewski PA, Garrison A, Wirtz AJ, Akguen S, Brandel-Ankrapp K, Endege W, Boyce FM, Ohnishi YN, Mazei-Robison M, Maze I, Neve RL, Robison AJ (2020) Circuit-specific hippocampal ΔFosB underlies resilience to stress-induced social avoidance. Nat Commun 11:4484.

Erwin SR, Sun W, Copeland M, Lindo S, Spruston N, Cembrowski MS (2020) A Sparse, Spatially Biased Subtype of Mature Granule Cell Dominates Recruitment in Hippocampal-Associated Behaviors. Cell Rep 31:107551.

Fischer KB, Collins HK, Callaway EM (2019) Sources of off-target expression from recombinase-dependent AAV vectors and mitigation with cross-over insensitive ATG-out vectors. Proc Natl Acad Sci USA.

Gibb B, Gupta K, Ghosh K, Sharp R, Chen J, Van Duyne GD (2010) Requirements for catalysis in the Cre recombinase active site. Nucleic Acids Res 38:5817–5832.

Gong S, Doughty M, Harbaugh CR, Cummins A, Hatten ME, Heintz N, Gerfen CR (2007) Targeting Cre recombinase to specific neuron populations with bacterial artificial chromosome constructs. J Neurosci 27:9817–9823.

Guenthner CJ, Miyamichi K, Yang HH, Heller HC, Luo L (2013) Permanent genetic access to transiently active neurons via TRAP: targeted recombination in active populations. Neuron 78:773–784.

Hilton IB, D’Ippolito AM, Vockley CM, Thakore PI, Crawford GE, Reddy TE, Gersbach CA (2015) Epigenome editing by a CRISPR-Cas9-based acetyltransferase activates genes from promoters and enhancers. Nat Biotechnol 33:510–517.

Jinek M, Chylinski K, Fonfara I, Hauer M, Doudna JA, Charpentier E (2012) A programmable dual-RNA-guided DNA endonuclease in adaptive bacterial immunity. Science 337:816–821.

Kaspar BK, Vissel B, Bengoechea T, Crone S, Randolph-Moore L, Muller R, Brandon EP, Schaffer D, Verma IM, Lee K-F, Heinemann SF, Gage FH (2002) Adeno-associated virus effectively mediates conditional gene modification in the brain. Proc Natl Acad Sci USA 99:2320–2325.

Konermann S, Brigham MD, Trevino AE, Joung J, Abudayyeh OO, Barcena C, Hsu PD, Habib N, Gootenberg JS, Nishimasu H, Nureki O, Zhang F (2015) Genome-scale transcriptional activation by an engineered CRISPR-Cas9 complex. Nature 517:583–588.

Kühn R, Schwenk F, Aguet M, Rajewsky K (1995) Inducible gene targeting in mice. Science 269:1427–1429.

Kumar N, Stanford W, de Solis C, Aradhana, Abraham ND, Dao T-MJ, Thaseen S, Sairavi A, Gonzalez CU, Ploski JE (2018) The Development of an AAV-Based CRISPR SaCas9 Genome Editing System That Can Be Delivered to Neurons in vivo and Regulated via Doxycycline and Cre-Recombinase. Front Mol Neurosci 11:413.

Lee G, Saito I (1998) Role of nucleotide sequences of loxP spacer region in Cre-mediated recombination. Gene 216:55–65.

Lobo MK, Karsten SL, Gray M, Geschwind DH, Yang XW (2006) FACS-array profiling of striatal projection neuron subtypes in juvenile and adult mouse brains. Nat Neurosci 9:443–452.

Madisen L et al. (2012) A toolbox of Cre-dependent optogenetic transgenic mice for light-induced activation and silencing. Nat Neurosci 15:793–802.

Metzger D, Chambon P (2001) Site- and time-specific gene targeting in the mouse. Methods 24:71–80.

Mima K, Deguchi S, Yamauchi T (2001) Characterization of 5’ flanking region of alpha isoform of rat Ca2+/calmodulin-dependent protein kinase II gene and neuronal cell type specific promoter activity. Neurosci Lett 307:117–121.

Morelli AE, Larregina AT, Smith-Arica J, Dewey RA, Southgate TD, Ambar B, Fontana A, Castro MG, Lowenstein PR (1999) Neuronal and glial cell type-specific promoters within adenovirus recombinants restrict the expression of the apoptosis-inducing molecule Fas ligand to predetermined brain cell types, and abolish peripheral liver toxicity. J Gen Virol 80 (Pt 3):571–583.

Pickar-Oliver A, Gersbach CA (2019) The next generation of CRISPR-Cas technologies and applications. Nat Rev Mol Cell Biol 20:490–507.

Savell KE, Bach SV, Zipperly ME, Revanna JS, Goska NA, Tuscher JJ, Duke CG, Sultan FA, Burke JN, Williams D, Ianov L, Day JJ (2019a) A Neuron-Optimized CRISPR/dCas9 Activation System for Robust and Specific Gene Regulation. Eneuro 6.

Savell KE, Day JJ (2017) Applications of crispr/cas9 in the mammalian central nervous system. Yale J Biol Med 90:567–581.

Savell KE, Gallus NVN, Simon RC, Brown JA, Revanna JS, Osborn MK, Song EY, O’Malley JJ, Stackhouse CT, Norvil A, Gowher H, Sweatt JD, Day JJ (2016) Extra-coding RNAs regulate neuronal DNA methylation dynamics. Nat Commun 7:12091.

Savell KE, Sultan FA, Day JJ (2019b) A Novel Dual Lentiviral CRISPR-based Transcriptional Activation System for Gene Expression Regulation in Neurons. Bio Protoc 9.

Savell KE, Tuscher JJ, Zipperly ME, Duke CG, Phillips RA, Bauman AJ, Thukral S, Sultan FA, Goska NA, Ianov L, Day JJ (2020) A dopamine-induced gene expression signature regulates neuronal function and cocaine response. Sci Adv 6:eaba4221.

Savitt JM, Jang SS, Mu W, Dawson VL, Dawson TM (2005) Bcl-x is required for proper development of the mouse substantia nigra. J Neurosci 25:6721–6728.

Schnütgen F, Doerflinger N, Calléja C, Wendling O, Chambon P, Ghyselinck NB (2003) A directional strategy for monitoring Cre-mediated recombination at the cellular level in the mouse. Nat Biotechnol 21:562–565.

Shaul O (2017) How introns enhance gene expression. Int J Biochem Cell Biol 91:145–155.

Shechner DM, Hacisuleyman E, Younger ST, Rinn JL (2015) Multiplexable, locus-specific targeting of long RNAs with CRISPR-Display. Nat Methods 12:664–670.

Tsien JZ, Chen DF, Gerber D, Tom C, Mercer EH, Anderson DJ, Mayford M, Kandel ER, Tonegawa S (1996) Subregion- and cell type-restricted gene knockout in mouse brain. Cell 87:1317–1326.

Yeo NC et al. (2018) An enhanced CRISPR repressor for targeted mammalian gene regulation. Nat Methods 15:611–616.

Zhang J, Zhang L, Jiao H, Zhang Q, Zhang D, Lou D, Katz JL, Xu M (2006) c-Fos facilitates the acquisition and extinction of cocaine-induced persistent changes. J Neurosci 26:13287–13296.

Zhuang X, Masson J, Gingrich JA, Rayport S, Hen R (2005) Targeted gene expression in dopamine and serotonin neurons of the mouse brain. J Neurosci Methods 143:27–32.

